# Challenging age-structured and first order transition cell cycle models of cell proliferation

**DOI:** 10.1101/2023.09.08.556865

**Authors:** Paolo Ubezio

## Abstract

Uncontrolled cell proliferation is the key feature of tumours. Because experimental measures provide only a partial view to the underlying proliferative processes, such as cell cycling, cell quiescence and cell death, mathematical modelling aims to provide a unifying view of the data with a quantitative description of the contributing basic processes. Modelling approaches to proliferation of cell populations can be divided in two main categories: those based on first order transitions between successive compartments and those including a structure of the cells’ life cycle. Here we challenge basic models belonging to the two categories to fit time course data sets, from our laboratory experience, obtained observing the proliferative phenomenon with different experimental techniques in a cancer cell line. We disclose the limitations of too simple models. At the minimal complexity level accounting for all available data the two approaches converge and suggest similar scenarios for the underlying proliferation process, in both untreated conditions and after treatment.

## Introduction

Cell proliferation is driven by cell cycling processes through G_1_, S, G_2_ and M phases, by the quiescence-proliferation interplay and by cell death processes. The increase of the number of individuals of a cell population originates from the cell cycle, where each cell progressively increases its mass, doubles its DNA content in phase S and finally divides in mitosis (M), giving birth to two daughter cells. This expansive mechanism is counterbalanced by quiescence and death processes, arresting cell cycle progression in a G_0_ phase, usually at the beginning of the cycle, where cells can stop for an indefinite time or be committed to die.

Cell cycle progression, at the cell population level, has been modelled since the sixties of the last century with balance equations [4, 12, 23], either assuming that the progression occurs through subsequent compartments or phases and the transit from a compartment to the next is a Poisson process (“first order transition” models), or using structured models, with structures based on cell age [25, 26], “maturation” [14], size [11], DNA content [2, 5]. First order transition models are described by a set of ordinary differential equations or even a single equation as in the core of most PK/PD models, where a cell population is represented with a single compartment. Structured models are instead described by partial differential equations like the Von Foerster equation [25] for an age-structured population, taking advantage of the fact that age increases exactly with time. The corresponding equations with other structures have a limited use because they include the rate at which cells increase the structuring variable, which is generally unknown and is expected to be highly variable among cells. The models are usually implemented in their discrete-time form with difference equations, for computing convenience and because the meaningful variations of the state variables, such as numbers of cells, typically occur over hours. Thus, difference equations with a not too short time step go possibly closer to the real world than differential equations, which treat the cell number or cell “density” as continuous variables subject to infinitesimal variations.

Although the adoption of simple models may be justified by lack of experimental data available to the modeler, excessive simplification may lead to unrealistic or wrong deductions. Several experimental methods can be used to obtain information on the kinetics of progression within the cell cycle, and rich datasets exist, particularly for tumour cell proliferation and the effects of treatment, that can be used to challenge the mathematical models. A direct comparison of the different modelling approaches has not been made before, in our knowledge, and is pursued in the present work, exploiting one of the databases previously published by our laboratory, and exploring the predictions and limitations of models at increasing complexity levels.

### The database

The dataset used here comes from our studies in vitro on the effects of anticancer treatments on the ovarian cancer cell line IGROV-1 [15, 16, 18, 19, 31], mainly from the work on the effects of X-rays published in Plos Computational Biology, where the details of the experimental methods are reported [13]. We will consider first untreated cells, which were in exponential phase at the beginning of the experiment (t=0), and then examples of treatment modelling. The IGROV-1 cell population was observed with independent experimental platforms, measuring: 1) cell cycle percentages by DNA flow cytometry (DNA_FC) (pooling G_2_ and mitotic cells as G_2_M), 2) the genealogical trees of individual cells by time lapse microscopy (TLM) and 3) absolute numbers of cells by coulter counter (ACC).

Main data are time courses up to t=72h or 96h of:

- the overall cell number (N(t))
- the percent of cells in G_1_, S and G_2_M cell cycle phases (%G_1_(t), %S(t), %G_2_M(t)),
- the percent of “generation 0” (gen0) cells, i.e. cells observed at t=0 and tracked until they divide or die (%Ngen0(t)), and of those in subsequent generations: %Ngen1(t), i.e. siblings of gen0 cells, %Ngen2(t), %Ngen3(t), etc.

Moreover, the intermitotic time of thousands of cells was individually directly recorded by TLM, giving the frequency distribution of the cell cycle time (F(T_c_)).

In addition, we considered short-time bi-parametric flow cytometric measures of a pulse-chase experiment with bromodeoxyuridine (BrdU), where the cell cycle progression of cells that were in S-phase at t=0 (keeping the BrdU label and thus named BrdU+ cells) was followed up to 12h, giving the overall percentage of BrdU+ cells (%BrdU+(t)) and the percentage of residual undivided BrdU+ cells (%Res+(t)).

## Results

Aim of the present study is a conceptual comparison of the capability of the first order transition and age-structured models to reproduce the observations. We will challenge the models with the data of IGROV-1 cells, to obtain for each model a set of best-fit parameter values for the unperturbed growth of this tumour cell line and then for X-rays treatments.

### The growth curve and the steady state of balanced growth

The experimental growth curve of an untreated cell population of IGROV-1 cells was exponential in the first 48h with negligible cell death, and thus easily fitted also by a simple one-compartment first order model without consideration of cell cycle phases (Model A, Fig. 1). Subsequent decrease of growth rate due to reduction of nutrients/space availability could be considered introducing a time dependence of parameters, or using a traditional Gompertz equation, but we will suppose for the moment to limit our observation to a time period where the growth curve is exponential with good approximation.

**Fig. 1.**
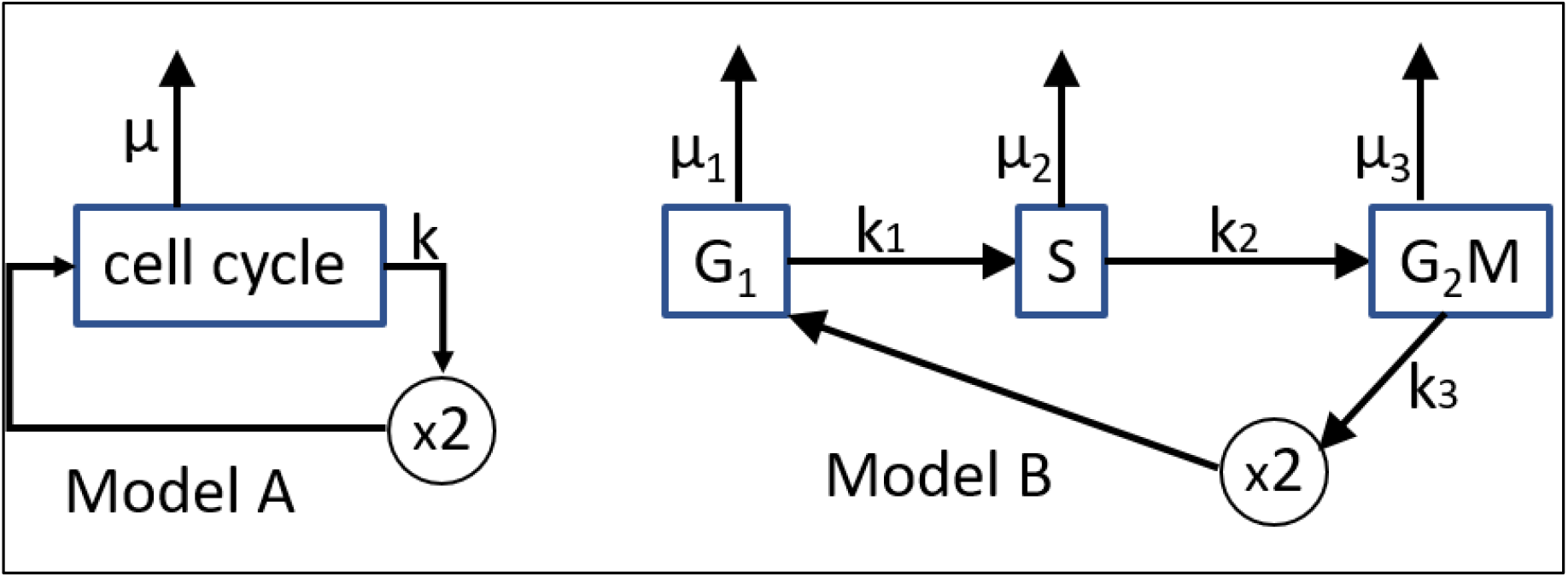
Model A. One-compartment first order model: dN(t)/dt =(*k*-*μ*)·N(t), where *k* is the proliferation rate and *μ* the death rate. Model B. Model with first order transitions between cell cycle phases: dN_G1_/dt=2*k*_*3*_·N_G2M_-(*k*_*1*_+*μ*_*1*_)·N_G1_; dN_S_/dt=*k*_*1*_·N_G1_-(*k*_*2*_+*μ*_*2*_)·N_S_; dN_G2M_/dt=*k*_*2*_·N_S_-(*k*_*3*_+*μ*_*3*_)·N_G2M_ where *k*_*1*_, *k*_*2*_, *k*_*3*_ are exit rates from G_1_, S and G_2_M respectively and *μ*_*1*_, *μ*_*2*_, *μ*_*3*_ the corresponding death rates. The state variables N_G1_(t), N_S_(t) and N_G2M_(t) are the number of cells in cell cycle phases, directly related to the observable quantities: N(t)=N_G1_(t)+N_S_(t)+N_G2M_(t), %G_1_(t)=100·N_G1_(t) /N(t), %S(t)=100·N_S_(t) /N(t) and %G_2_M(t)=100·N_G2M_(t) /N(t)

Obviously Model A allows to deduce the “growth rate” (*α* =*k*-*μ*) from the growth curve alone, while would require an independent measure of the death rate *μ* in order to specify the proliferation rate *k*. Because there are several methods to evaluate cell death, one may expect that the death rates could be directly measured. Unfortunately, this is not the case. The popular methods are based on probes of cell death, detecting the cells which are in a window of the dying process, thus giving the percent of cells in that time-window (of unspecified width) at the particular time of the measure. The relationship of this percentage with the death rate or the overall number of dead cells in a time interval remains elusive. That’s why a proper use of these markers is limited to a qualitative evaluation of the presence of death. In our experiments we considered cell death negligible when measuring < 3% positive cells detected by one of these markers. Nevertheless, the information of presence/absence of cell death is important to steer the choice of considering or not the death parameter *μ*.

Although model A is often adopted in proliferation modelling, particularly when only the overall cell number or mass is available (e.g. in the PK/PD field), its description of the phenomenon is clearly too simplistic. It is enough to consider a simple measure of cell cycle percentages %G_1_, %S %G_2_M, to be forced to abandon model A for a more complex one, such as model B, including consideration of G_1_, S and G_2_M phases.

Coupling measures of the growth curve with cell cycle percentages imposes a strong constrain, which allows to deal several situations with model B avoiding over-parametrization, and in particular to make a deeper analysis of the exponential growth. In experimental research a balanced steady state of exponential growth is usually (often implicitly) assumed when measuring the characteristic properties of a cell line and it is the starting point in most experiments on effects of treatments.

Model B fits properly contemporaneously the number of cells and cell cycle percentages of the IGROV-1 cell line in the exponential growth (Fig. 2), as can be expected because, assuming the steady state and uniform death rate (*μ*=*μ*_*1*_*=μ*_*2*_*=μ*_*3*_), the equations can be solved knowing the growth rate (directly calculated from the growth curve) and the constant values of %G_1_, %S and %G_2_M.

**Fig. 2.**
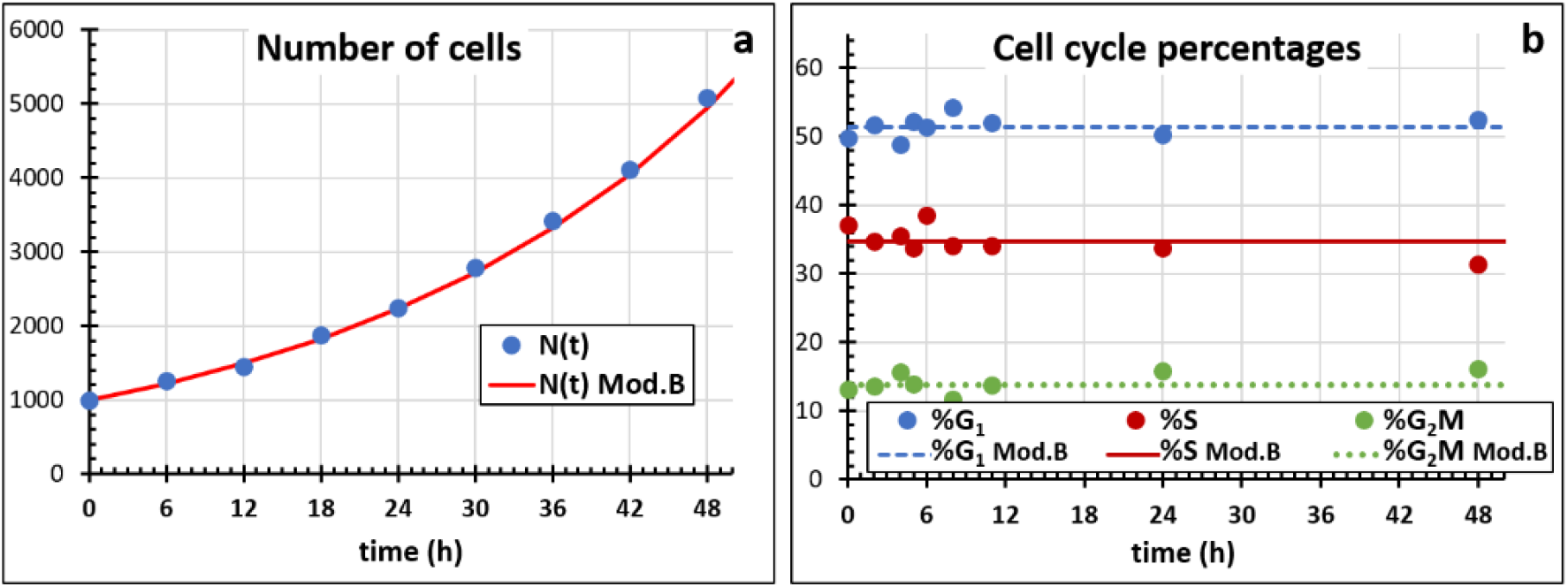
Contemporary fit of the number of cells N(t) (panel **a**) and cell cycle percentages %G1(t), S(t) and G2M(t) (panel **b**) in the exponential phase of growth with Model B with three parameters: *k*_*1*_, *k*_*2*_, *k*_*3*_, (with fixed death rates: *μ*_*1*_*= μ*_*2*_*= μ*_*3*_=0), assuming the desynchronised starting distribution associated with parameters’ values. (N(t) normalised to N(0)=1000)

It is interesting to consider briefly the desynchronization process, which is also used in the fitting procedure at each variation of the parameters to achieve the starting cell cycle distribution. Fig. 3a shows how model B realizes the desynchronization to the unique balanced steady state for the IGROV-1 cell line from different starting distributions. Even assuming that at a given time the cells are all in a single cell cycle phase, the model B desynchronizes the cell population in a very short time (variation of cell cycle percentages become less than 1% in 20 hours, less than one doubling time). This can be compared with published data on the same cell line [10] for cells initially in phase S (Fig. 3b). Data show a much slower desynchronization rate than predicted with model B, suggesting that the model does not catch properly the actual flow of the cells through cell cycle phases. In fact, according to models A and B, cells can complete the whole cell cycle in an infinitesimal time, while in reality it takes at least hours only for DNA replication and mitosis.

**Fig. 3.**
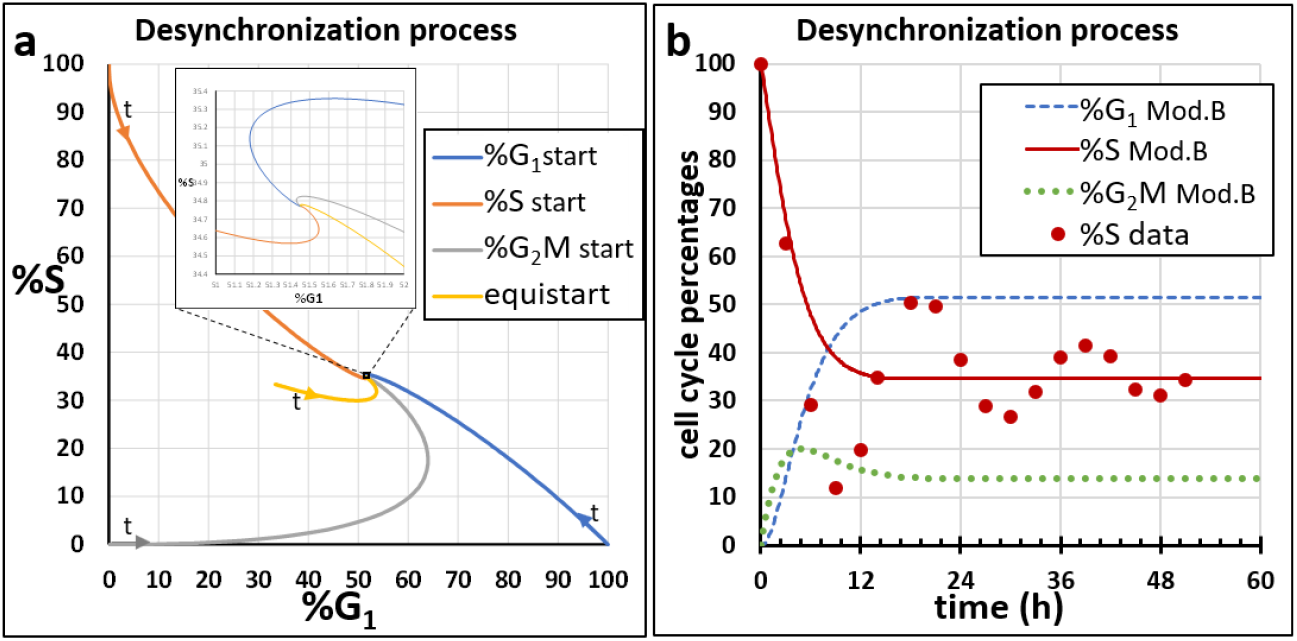
The desynchronization process in IGROV-1 cells as simulated by the best fit model B with parameters *k*_*1*_, *k*_*2*_, *k*_*3*_ as in Fig. 2. **a** Approach to the unique steady state starting with 100% cells in G_1_ (“G1 start”), in S (“S start”), in G_2_M (“G2M start”) or 33.33% in each phase (“equistart”), in a %S vs %G1 plot with time as parameter of the curves [25]. **b** Time course of cell cycle percentages starting with 100% S phase cells, as simulated by the model (“%G_1_ Mod.B”, “%S Mod.B” and “%G_2_M Mod.B”) compared with experimental data (“%S”) [10]

In the case of IGROV-1 cells in the exponential growth phase, independent experiments demonstrated that both quiescence and cell death were negligible, so it was acceptable to assume *μ*_*1*_*= μ*_*2*_*= μ*_*3*_=0 and not to consider quiescence, but in general this is not true, particularly *in vivo*.

Model C (Fig. 4) is a minimal model including all three pillars concurring to proliferation, adding a quiescence (“G_0_”) compartment [14] to the model B, which already considered cell cycle and cell death (or, more generally, “cell loss”). In this model, newborn cells may remain proliferating with probability θ or are committed to become quiescent (probability 1-θ); then quiescent cells die with rate *μ*_*0*_ or re-enter the cell cycle, with rate *k*_*0*_.

**Fig. 4.**
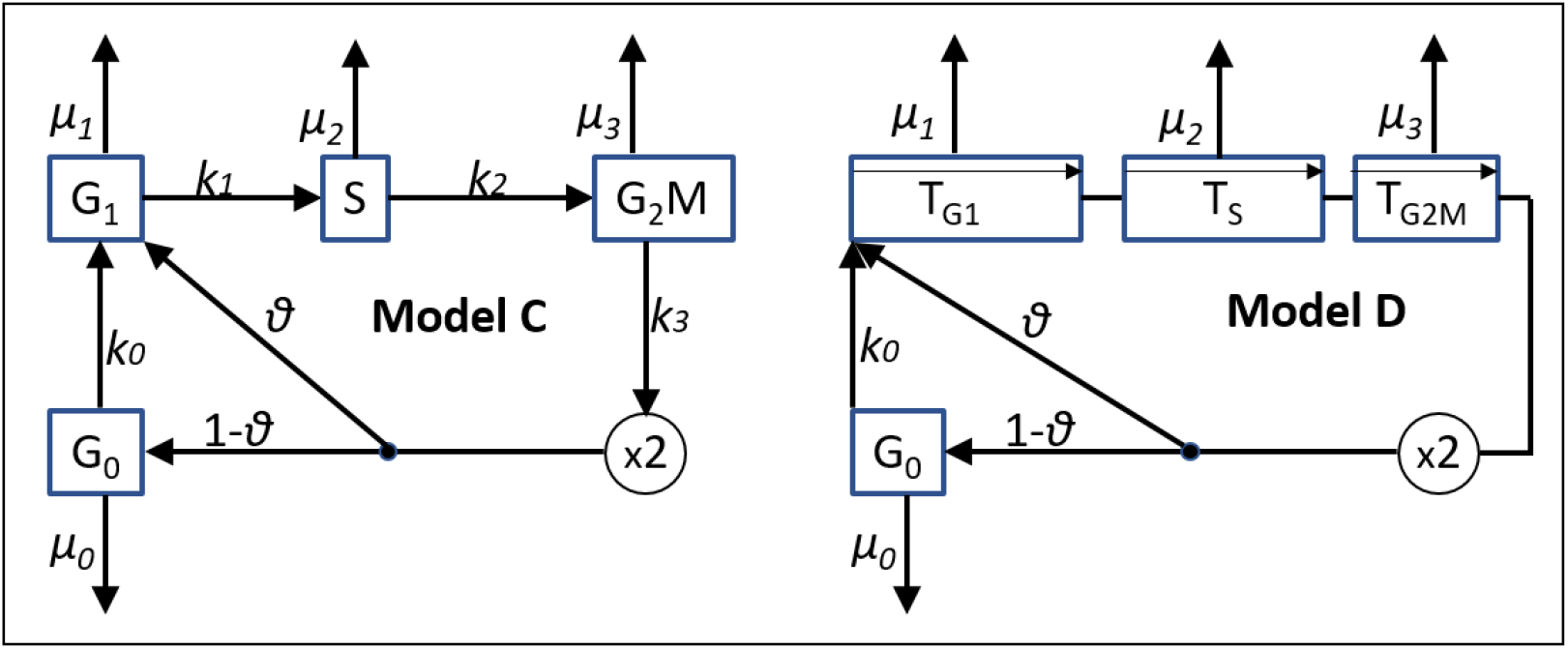
Models including the quiescence compartment. Model C: first order transition model. Model D: Bertuzzi-Gandolfi age-structured model. In both models, quiescence is rendered with a similar G_0_ compartment. The parameters of the G_0_ compartment are: *θ* (fraction of cells bypassing G_0_), *k*_*0*_ (rate of re-entering in cycle of quiescent cells), *μ*_*0*_ (rate of death of quiescent cells). G_1_, S G_**2**_M parameters are phase exit rates *(k*_*1*_, *k*_*2*_, *k*_*3*_) in model C, phase durations (*T*_*G1*_, *T*_*S*_, *T*_*G2M*_) in model D, phase death rates (*μ*_*1*_, *μ*_*2*_, *μ*_*3*_). Simpler models without G_0_, as model B, are obtained by setting *θ*=1

Similarly, model D (Bertuzzi-Gandolfi model [6]) is a basic age-structured model with fixed phase durations *T*_*G1*_, *T*_*S*_, *T*_*G2M*_. In model D all cells entering G_1_ at t=t’ will exit G_1_ at t=t’+T_G1_, if they do not die in the meantime, and similarly for S and G_2_M phases. Within each phase, the ages *a*_*G1*,_ *a*_*S*,_ *a*_*G2M*_ are measured from cell entry in the corresponding phase. In the discrete-time formulation, each phase is divided in small age intervals Δ*a* equal to the step time Δt. Considering that the cells of age between *a*_*G1*_ and *a*_*G1*_+Δ*a* (with 0 ≤*a*_*G1*_ <T_G1_) at time *t* are those in the previous age interval in the previous step time which do not die in Δ*t*, the simple balance equation holds: N_G1_(*a*_*G1*_, *t*) = (1-*μ*_*G1*_·Δ*t*)·N_G1_(*a*_*G1*_-Δ*a, t*-Δ*t*). Cells exiting G_1_ enter S at *a*_*S*_=0 while cells entering at *a*_*G1*_=0 come from G_2_M exit and division. S and G_2_M age structures act in the same way, while G_0_ is modelled as a single compartment, with first-order exit rate (*k*_*0*_) and without age structure, as in model C.

The introduction of G_0_ makes cell cycle models flexible to account for different biological hypotheses. For instance, it is possible to consider the case in which all cells necessarily pass through G_0_ by setting θ=0 and *k*_*0*_>0. Conversely, with θ>0 and *k*_*0*_=0 models C and D interpret G_0_ as a compartment of definitively quiescent cells out of cycle. Moreover, G_0_ compartment can be viewed as a compartment of quiescent stem cells, recruited to proliferation with rate *k*_*0*_.

It should point out that a univocal balanced steady state of exponential growth exists also in these and in more complex models derived from them, as far as parameters are constant in time (see reviews [1, 33] of the mathematical literature on this topic). A variation of some of the parameters, possibly caused by variation of environmental factors, would lead the population towards a different steady state. As an example of this phenomenon we will consider in the next section the approach to confluence.

### The approach to confluence

The steady state of exponential growth cannot be maintained indefinitely, both *in vitro* and *in vivo*, because nutrients are progressively consumed and/or the available space reduces. In this case we typically observe a slowdown of the proliferation with an increase of %G1, indicating that at least a fraction of the cells where delayed or arrested before the onset of DNA synthesis. This motivated the introduction in the models of a compartment (G_0_) collecting cells committed to become quiescent before/within G_1_. From the experimental point of view, if cell cycle percentages are calculated based on DNA cellular content only, proliferating (G_1_) and quiescent (G_0_) cells are indistinguishable and “%G_1_” is renamed as “%G_0_G_1_”. Fig. 5 shows how models C and D fit the data of the IGROV-1 cell population in an extended 0-96h interval, characterized by an increase of %G_0_G_1_, a decline of the growth rate and detection of apoptotic cells at 96h.

**Fig. 5.**
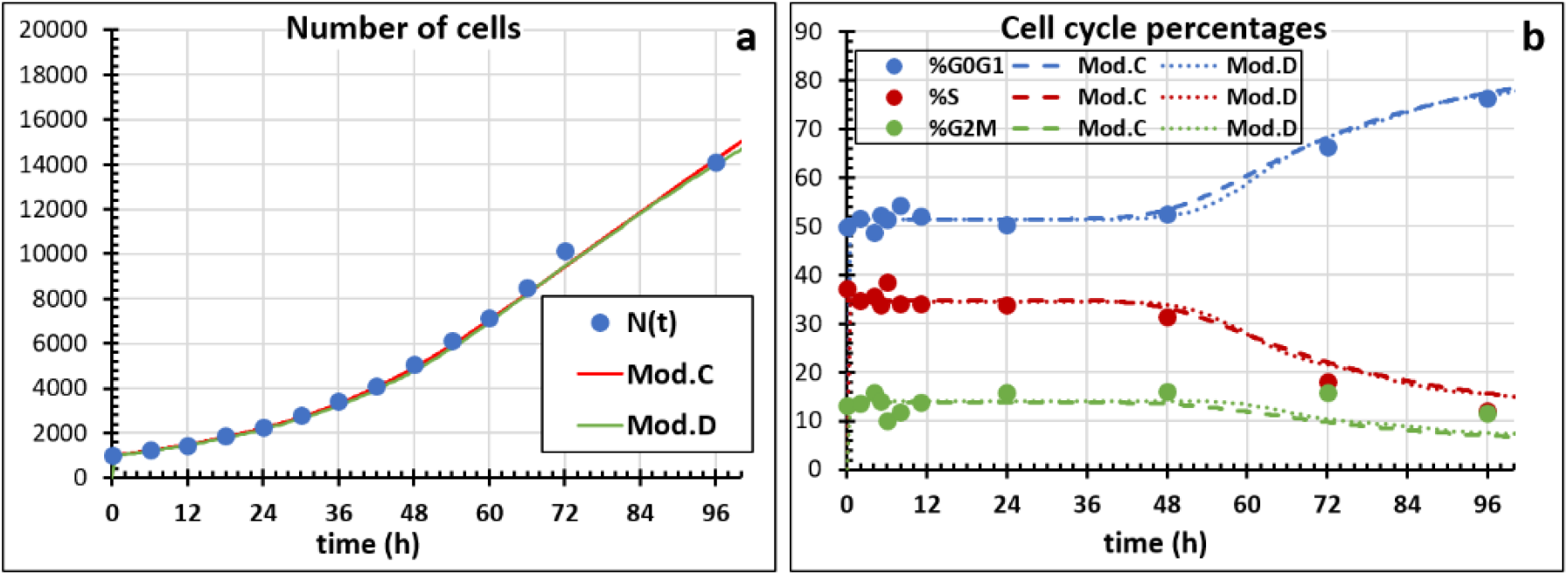
Contemporary fit of the number of cells N(t) (panel **a**) and cell cycle percentages %G_0_G_1_(t), S(t) and G_2_M(t) (panel **b**) in the 0-96h interval, with Models C (dashed lines) and D (dotted lines). Respective steady state G_1_, S G_**2**_M parameters were fixed (obtained from a previous steady state fit in 0-48h) and the approach to confluence was fitted varying parameters *θ* and *μ*_*0*_ in the 48-96h interval

In order to fit the whole 0-96h time course, the steady state best fit G_1_, S, G_**2**_M parameters were applied in the 0-48h interval (as shown in Fig. 2 for model C and similarly for model D), then two additional parameters were made variable in the 48-96h interval only: *θ*, enabling cells to enter G_0_, and *μ*_*0*_, accounting for G_0_ cell loss (*θ*=1 and *μ*_*0*_ =0 in the 0-48h interval). The best fits shown in Fig. 5, obtained optimizing *θ* and *μ*_*0*_, are satisfactory for both types of models, although somewhat underestimating %G_2_M at 72 and 96h. This suggests the presence of quiescent cells also in G_2_, that could be accounted introducing an additional compartment for them. We conclude that time courses of the number of cells and cell cycle percentages are not sensitive to discriminate between the two types of models.

### More flow cytometric experiments: BrdU labeling

In order to challenge/identify the models let’s consider additional experiments with the FC platform. The BrdU labeling method provides valuable data for this purpose. The method can be applied both *in vitro* and *in vivo* and its use to evaluate the potential doubling time of tumours in patients was widespread in the last two decades of the last century [3]. It is based on the labeling of S phase cells with BrdU, which are then followed in time while exiting S, entering G_2_M, divide and enter G_1_ for a new cycle. Usually these BrdU+ cells remain distinguishable for at least one cell cycle time, providing data suitable for cell cycle modelling or testing consistency of proposed models. On the other hand, manipulation *in vitro* of the cells for BrdU labeling may cause a temporary cell cycle delay, which may be considered to refine models, and identification of the subpopulation of BrdU+ may be challenging in some instances.

Fig. 6 shows the time courses of two kind of data obtained in a pulse labeling BrdU experiment: the percentage of labelled cells (%Brdu+(t)) and of residual undivided labelled cells (%ResU+(t)). The percentage of labelled cells (%BrdU+(t)) includes all BrdU+ cells at time “t”, i.e. those still undivided and their descendant, while the percentage of residual undivided labelled cells (%ResU+(t)) refers to undivided cells among those that were in S phase at t=0. Notice that such data can be readily obtained by simulation with the above models: given a set of parameter values and obtained the initial cell cycle distribution as before, the simulation of the time course will be made two times, one for BrdU+ cells (starting with S phase cells only, which have been labelled at t=0), and one for BrdU-cells (starting with G_1_ and G_2_M phase cells only, which remain unlabelled). The time course of the whole population is trivially obtained by summing the number of BrdU+ and BrdU-in each phase at each time. Refinement of the modelling may include consideration of the finite duration of the “pulse” (exposure to BrdU lasts 15-30min, i.e. 1-2% of a typical cell cycle time).

**Fig. 6.**
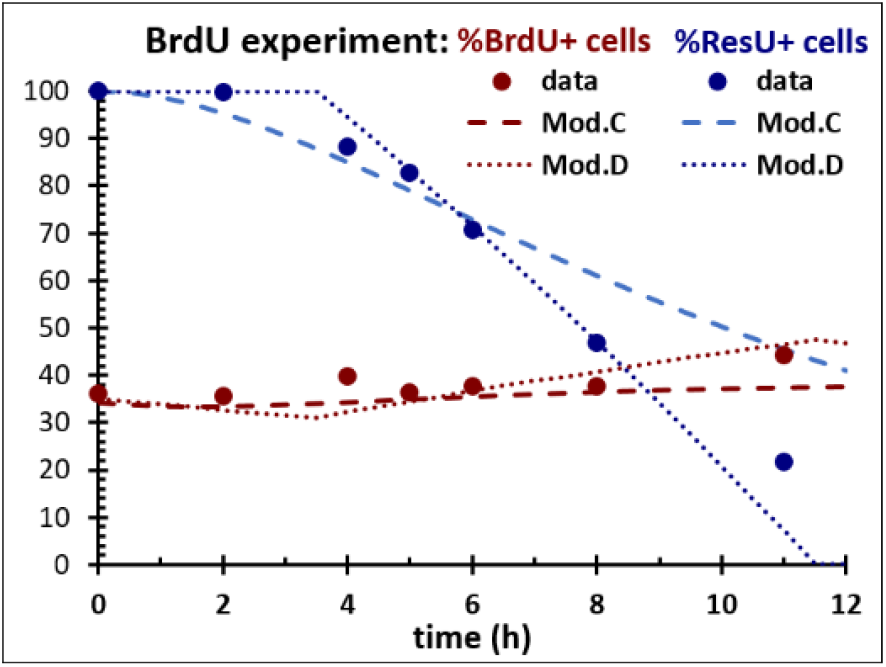
BrdU labeling experiment in IGROV-1 cells: time course of the percentage of labelled cells (%BrdU+) and of residual undivided labelled cells (%ResU+). Model C (dashed lines) and D (dotted lines) predictions with the best fit parameters obtained by contemporary fit of N(t) and %G1(t), S(t) and G2M(t) in the exponential phase of growth

Fig. 6 evidences the difficulties of the model C to reproduce the time course of %ResU+, while model D shows the expected two periods of %ResU+(t): the first one when no division was observed (%ResU+ =100%) corresponding to the time to traverse G_2_M phase before dividing, and the second period characterised by a progressive decrease of %ResU+, as the cells gradually arrive in mitosis and divide, leaving the first cycle. The bending of the data respect to the sharp division of the two periods predicted by model D, demonstrates that the duration of S and G2M is not the same for all cells, as assumed by model D.

### More experiments: the distribution of intermitotic times and generation wise data

Of course, variability of phase durations is expected, even in a situation of uniform exposure to nutrients and unrestricted space, as in our experiment with IGROV-1 cells *in vitro*. The cellular content of any protein at a given time in a given cell cycle point should be variable because it is the result of millions of intertwined molecular reactions occurring from the birth in each cell, which also starts with a variable mass from the mitosis of the mother cell. Experimentally, the variability of cell cycle time can be directly measured by time-lapse microscopy (or “microcinematography”). In a pioneering work in 1963 from our laboratory [20], researchers applied microcinematography to measure the intermitotic times of individual cells by recording the time between two successive divisions. This experiment and several similar ones demonstrated that a minimum time for completing the cell cycle is necessary and variability was high with a frequency distribution skewed on the right. To account for that, Smith and Martin and others [7, 21] proposed a model (model SM) with two phases: the first (named “A-state”) with first order exit kinetics acting like the G_1_ phase of model B (but actually covering only a part of G_1_) followed by the second phase with fixed duration, covering the rest of the cell cycle. Model SM is equivalent to model D with *θ*=0, *k*_*0*_ > 0 and *μ*_*0*_ =0. In the Smith-Martin’s view the stochastic exit from the A-state was introduced to explain the observed inter-cell variability of the cell cycle times and desynchronization. However also model SM, as the simpler models A, B, C described above, fails to reproduce a typical frequency distribution of Tc (F(Tc)) (Fig. 7) as that observed in the IGROV-1 cell line, obtained with a modern equipment for time lapse microscopy (TLM).

**Fig. 7.**
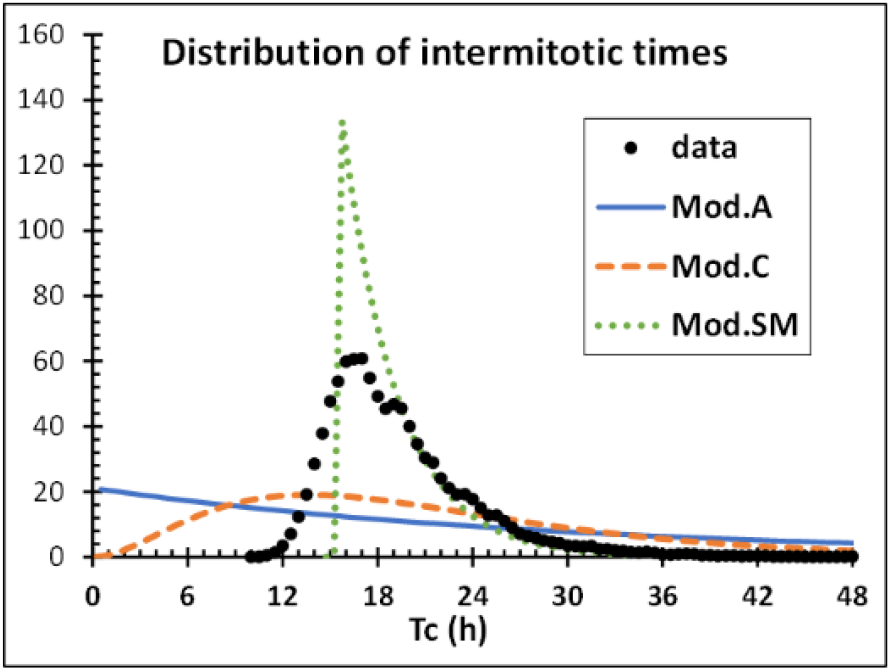
Frequency distribution of cell cycle times (F(Tc)) measured with TLM in IGROV-1 cells with 0.5 h intervals, normalized to 1000 cells and fitted with models A, C and SM

In the effort to reproduce this observation within the first order transition approach, model C can be modified splitting each phase in sub-compartments, characterised by the same exit rate [24]. In this way, the duration of each phase comes to follow an Erlang distribution and model parameters become the exit rates from sub-compartments (*k*_*1*_ for G_1_ sub-compartments, *k*_*2*_ for S, *k*_*3*_ for G_2_M) and the number of sub-compartments (*n*_*1*_, *n*_*2*_, *n*_*3*_) in each phase (model C_E). Instead the age structure of model D makes it suitable for the implementation of any distribution of phase durations. We implemented different two-parameter distributions [30], keeping the mean and coefficient of variation of phase durations as model parameters: *T*_*G1*_, *T*_*S*_, *T*_*G2M*_, *CV*_*TG1*_, *CV*_*S*_, *CV*_*TG2M*_. Already Sisken and Morasca observed also that 1/Tc was normally distributed [20] and a thorough examination of several similar datasets by Castor demonstrated that best fits were obtained assuming reciprocal-normal distributions [8]. We will consider an extension of model D with reciprocal-normal distributions of phase durations (model D_R). Fig. 8a shows that models C_E and especially D_R provide a good fit of F(Tc).

**Fig. 8.**
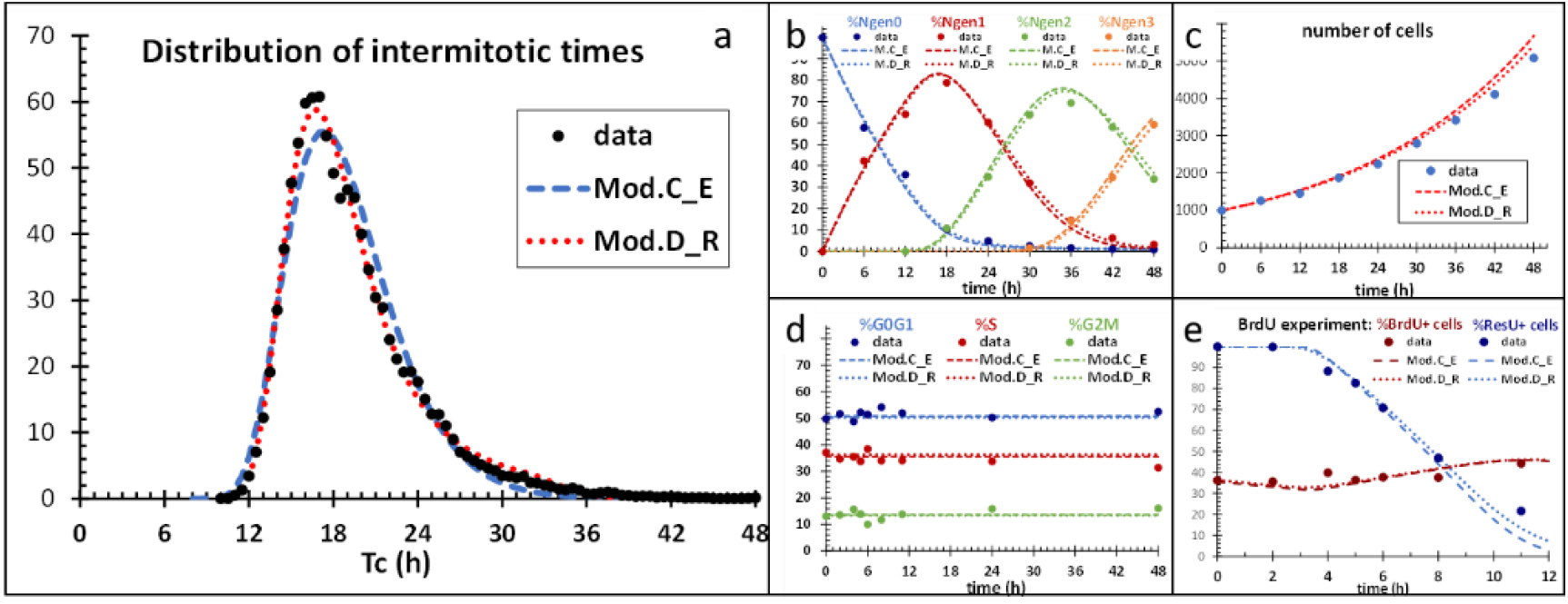
Model C_E and D_R predictions with the best fit parameters obtained by contemporary fit of all data of IGROV-1 cells in the exponential phase of growth: **a** frequency distribution of cell cycle times (TLM); **b** percent of cells in subsequent generations (TLM); **c** absolute cell count; **d** %G1(t), S(t), G2M(t) (FC); **e** %BrdU+, %ResU (FC). Model C_R parameters: *k*_*1*_, *k*_*2*_, *k*_*3*_, *n*_*1*_, *n*_*2*_, *n*_*3*_; model D_R parameters: *T*_*G1*_, *T*_*S*_, *T*_*G2M*_, *CV*_*TG1*_, *CV*_*S*_, *CV*_*TG2M*_

Using TLM it is also possible to measure directly the expansion of the cell population in a generation-wise perspective [13]. This is achieved by tracking each cell detected at the start of the experiment in several fields of view and their descendants. In this way, it is possible to measure at each time the percentage of cells belonging to gen0, gen1 and successive generations.

Cell generation percentages for IGROV-1 cells in the exponential state are shown in Fig. 8b. In order to reproduce TLM data in the modelling framework, it is enough to replicate the cell cycle, so that the daughters of dividing cells instead of re-entering in the same cell cycle enter in a new cell cycle, with the same parameters. The state variables become *N*(*ph,a,gen,t*), giving the number of cells in phase “*ph*” (G_1_, S or G_2_M), generation “*gen*”, sub-compartment (model C-E) or age interval (model D_R) “*a*”, at time “*t*”, and *N*(*G*_*0*_,*gen,t*) for the number of G_0_ cells. For each cell ending G_2_M of generation “i”, two newborn cells enter at *a*_*G1*_=0 in generation “i+1”. For what concerns F(Tc), a parallel simulation of G_1_, S and G_2_M transit starting with all cells at *a*_*G1*_=0 gives the time course of the number of undivided cells, from which the cumulative of F(Tc) and F(Tc) itself are readily derived.

Fig. 8 shows the best fit achieved fitting all data together with models C_E and D_R. Both models provided satisfactory fit of the data, thus unifying the views of the measures in the five panels with a single reconstitution of the underlying cell cycling of the cell population.

The two modelling approaches differs in F(Tc), where the reciprocal-normal functions adopted in model D_R allow an accurate fit of the shape of the experimental F(Tc), not achievable in model C_E. In general, the flexibility for the choice of the type of the distribution of the age-structured model gives a clear advantage for modelling different situations, particularly when a wide intercell variability is present as in the case of most tumour cell populations. Model C_E is in fact equivalent to an age-structured model with Erlang distributions of phase durations but links this variability to the number of compartments within each phase, which is not practical for fitting purposes.

On the other hand, model C_E fits the other data in panels 8b to 8e no less well than model D_R, with similar values of mean and variance of G_1_, S and G_2_M duration. This demonstrates that these types of data are poorly sensitive to the highest moments of the distribution, a result that was confirmed by evaluating other distributions in the age-structured model (not shown).

### Modelling treatment

The models discussed in the previous section can be readily adapted for studying cell proliferation in different contexts. For instance, the generation structure can be adapted for simulation of differentiation chains, as in studies of the lineages of the hematopoietic system, by reinterpreting generations as sequential differentiation levels and introducing a parameter of differentiation probability for the passage from a level to the next. In our laboratory, model D_R was adopted as basal representation of tumour proliferation in studies of the effects of various anticancer treatments. Treatment induces two kinds of perturbations of the basal proliferation process: cell cycle delay/arrest and cell kill. These are the functional end points of the cellular molecular reactions induced by the chemical/physical/biological agent (drug, radiation, antibody or others) used to treat the disease. Both delay/arrest and cell kill have time and dose/schedule dependence. Moreover, cells are diversely sensitive to treatment in the cell cycle phases and put in action different mechanisms of response in each phase, typically arresting cell cycle progression at checkpoints in G_1_ and G_2_M and reducing the rate of DNA replication in S. Then, arrested cells may repair the damage and resume cell cycle progression, but not rarely leaving some damage that lead to novel arrest and possibly death also in the subsequent generations. These complex phenomena can be modelled in the framework of the above models adding two minimal “checkpoint modules” (CpM) describing the dynamics of checkpoint arrest/resumption in G_1_ and G_2_M and a parameter rendering the reduction of the DNA synthesis rate in phase S [9, 15, 19]. Introducing such modules in model D_R we reproduced and explained the effects of X-ray treatment at increasing doses in the IGROV-1 cell line (see [13] for more details). We will show here how those data for a sublethal (0.5Gy) and a high (5Gy) dose are modelled with first order transition models, without (Fig. 9) and with (Fig. 10) introducing CpMs. The best fit of unperturbed growth considered above for each model provided the fixed parameters values of the basal proliferation and the time zero asynchronous cell cycle distribution.

**Fig. 9.**
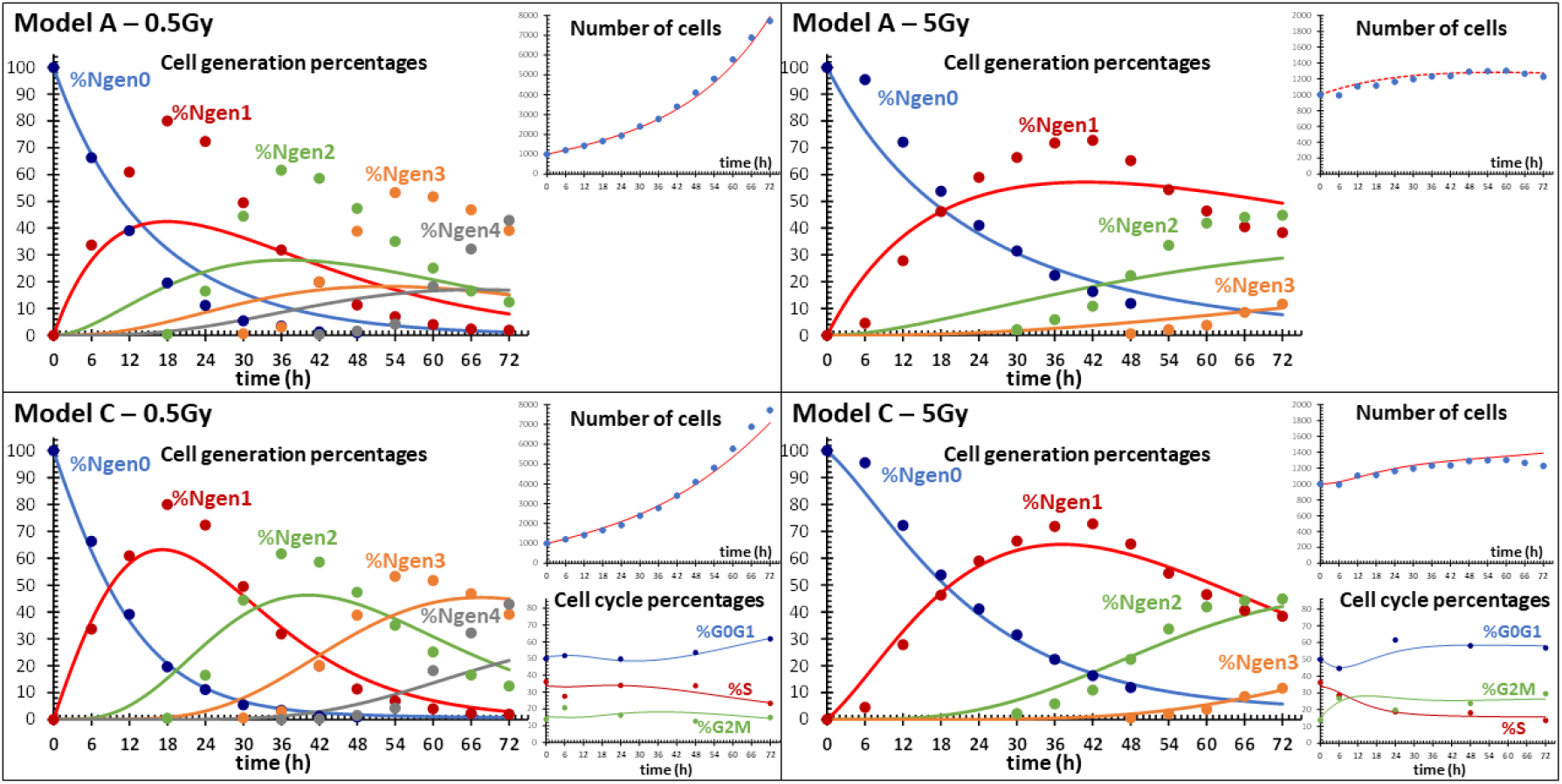
Treatment challenge. Data (circles) and best fit (lines) with models A (upper panels) and C (lower panels) of the time course of cell number within generations (time-lapse measures) and cell cycle phases (flow cytometry measures) after exposure to 0.5Gy (left panels) and 5Gy (right panels) of IGROV-1 cells *in vitro*. Treatment variables **Model A 0**.**5Gy:** *k*_*gen1,2*_ (i.e. with *k*_*gen2*_ = *k*_*gen1*_), **5Gy:** *k*_*gen0*_, *k*_*gen1,2*_, μ_*gen0,1,2*_; **Model C 0**.**5Gy:** *k*_*1gen1,2,3*_, *k*_*2gen1,2,3*_, *k*_*3gen1,2,3*_, *θ*_*gen3,4,5*_, **5Gy:** *k*_*1gen0*_, *k*_*2gen0,1,2,3*_, *k*_*3gen0*,_ *k*_*3gen1,2,3*_, μ_1*gen0,1,2,3*_. The other parameters were kept fixed as previously determined in untreated conditions

**Fig. 10.**
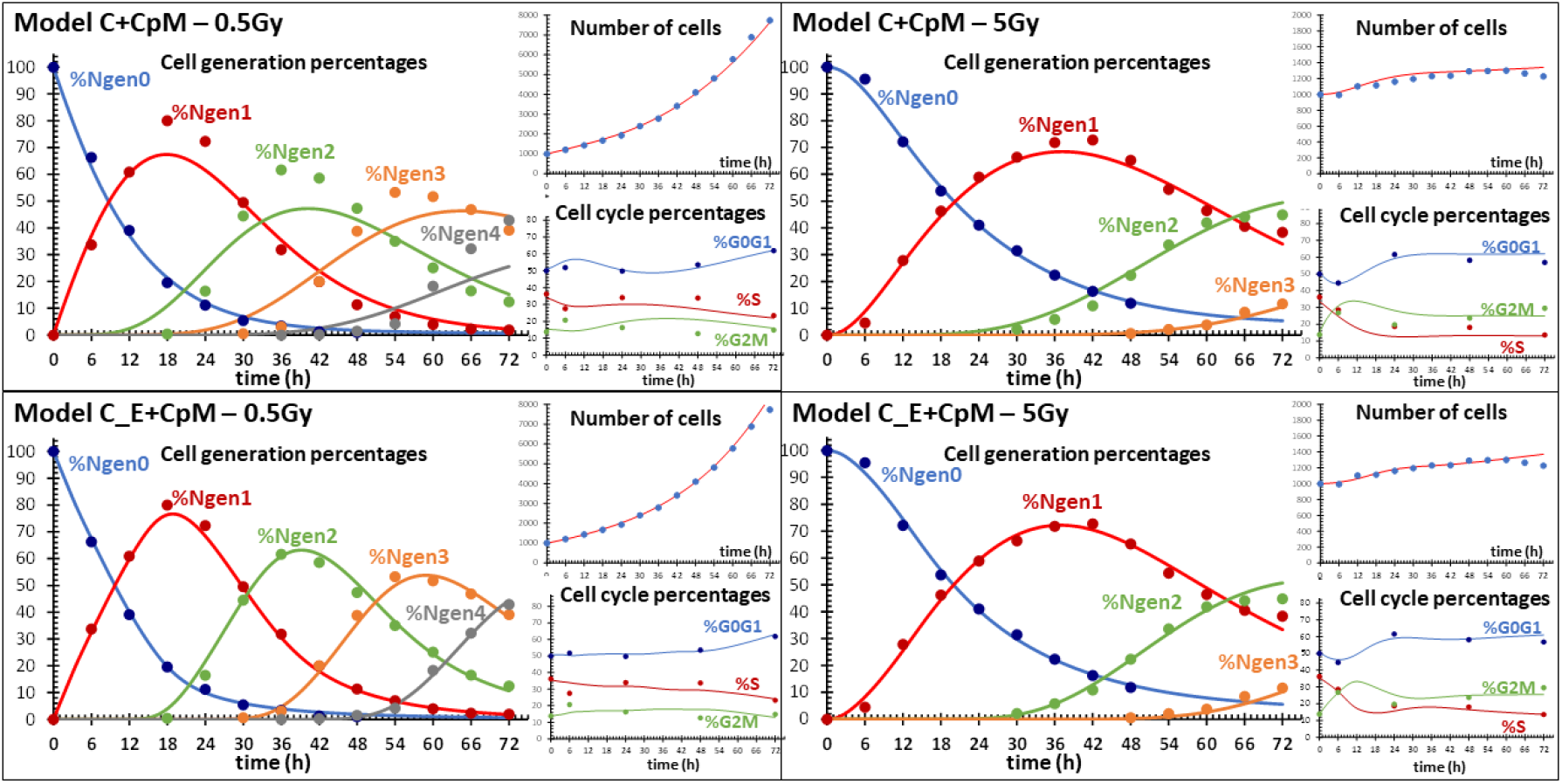
Treatment challenge. Data (circles) and best fit (lines) with models C+CpM (upper panels) and C_E+CpM (lower panels) of the time course of cell number within generations (time-lapse measures) and cell cycle phases (flow cytometry measures) after exposure to 0.5Gy (left panels) and 5Gy (right panels) of IGROV-1 cells *in vitro*. Treatment variables **Model C+CpM 0**.**5Gy:** *pBl*_*G1,gen1,2*_, *pBl*_*G2,gen0,1,2*_, *Rec*_*G1,gen1,2*_, *Rec*_*G2,gen0,1,2*_, *k*_*Sgen0,1*_, *θ*_*3,4,5*_, **5Gy:** *pBl*_*G1,gen0*_, *pBl*_*G1,gen1,2,3*_, *pBl*_*G1,gen0,1,2,3*_, *Rec*_*G1,gen0,1,2,3*_, *Rec*_*G2,gen0,1,2,3*_, *k*_*Sgen0,1*_, μ_G1*gen0,1,2,3*_, **Model C_E+CpM 0**.**5Gy:** *pBl*_*G1,gen0,1,2*_ (=*pBl*_*G2,gen0,1,2*_), *Rec*_*G1,gen1,2*_ (=*Rec*_*G2,gen0,1,2*_), *k*_*Sgen0,1*_, *θ*_*3,4,5*_, **5Gy:** *pBl*_*G1,gen0,1,2,3*_, *pBl*_*G2,gen0,1,2,3*_, *Rec*_*G1,gen0,1,2,3*_, *Rec*_*G2,gen0,1,2,3*_, *k*_*Sgen0,1*_, μ_G1*gen0,1,2,3*_. The other parameters were kept fixed as previously determined in untreated conditions

In models A, B and C without CpM cell cycle delay/arrest and cell kill were rendered by lowering the previously determined steady state values of the exit rates *k* (model A) or *k*_*1*_, *k*_*2*_, *k*_*3*_ (model C) and introducing non-zero values for the death rates *μ* (model A) or *μ*_*1*_, *μ*_*2*_, *μ*_*3*_ (model C) in the presence of cell kill. These variations may be different for gen0 cells (directly exposed to treatment) and their descendants or even be time-dependent. Let’s call *k*_*gen0*_ the *k* value in gen0, *k*_*gen*1_ the *k* value in gen1 and so on for all model parameters. Thus, a preliminary study is required to identify the relevant variables and the need to introduce a time-dependence for some parameter.

0.5Gy was a sublethal dose with a nearly exponential N(t), with a lower growth rate respect to control. The time course of the number of cells was well caught also by the simple model A with a single variable *k*_*gen1,2*_ (with *k*_*gen2*_ = *k*_*gen1*_) and leaving unperturbed *k* in gen0, gen3 and following (Fig. 9 upper left panel). However, the insight into the generation percentages revealed the inconsistence of that model, which predicted a too high number of cells remaining in gen0 after 6h and anticipated by hours the onset of cells in gen2 and subsequent generations. With the cytotoxic 5Gy dose, a good fitting of N(t) was obtained with model A, using a death parameter *μ*_*0,1,2,3*_ and two distinct *k* for gen0 and gen1,2,3, but the model again failed to predict correctly the passage through the subsequent generations (Fig. 9 upper right panel).

Models B and C reduced but did not eliminate the inconsistences of model A. Fitting 0.5Gy data with model C required to lower the exit rates in all cell cycle phases in gen1, gen2 and gen3, demonstrating the presence of delays in all phases, stronger in G_2_M than in G_1_ and S (Fig. 9 lower left panel). Again, no effects were suggested for gen0 cells. Parameter *θ*, unrelated to treatment, was included to render the approach to confluence, observed in 0.5Gy as in control. Best fit of 5Gy data required adding a cell death parameter in G_1_ and a gen0 delay in all phases as well as reinforcing delays in gen1 and subsequent generations (Fig. 9 lower right panel).

Fig. 10 shows the fits of models C and C_E with CpM modules in G_1_ and G_2_M, while S-phase delay, when required, was again modelled by reduction of the exit rate *k*_*S*_. G_1_ and G_2_M CpMs are single compartments collecting arrested cells in the respective phase. The dynamics of arrest, repair and kill is reproduced with three parameters: the probability of enter the block (*pBl*), the recycling (*Rec*) and death (*μ*) rates. *Rec* and *μ* orient the fate of arrested cells, which may continue cycling in the next phase or die.

C+CpM model (Fig. 10 upper panels) provided a closer fit to %gen data respect to model C shown in Fig. 9, but cell cycle percentages were fitted less well, particularly at 0.5Gy. Best C+CpM model of 0.5Gy data required G_1_ and G_2_M blocks from gen1 and no S-phase effects, while 5Gy data required to extend blocks to gen0. Recycling rates from blocks were higher in G_2_M respect to G_1_ blocked cells and lower in 5Gy respect to 0.5Gy. Death was limited to 5Gy G_1_ blocked cells with a rate similar to the G_1_ recycling rate. Overall, the results suggest that C+CpM fitted better and offered a more consistent interpretation of the data than model C at cytotoxic doses, while model C performed somewhat better at low doses.

C_E+CpM model overcame the limitations of both C and C+CpM models and enabled the best fits (Fig.10 lower panels). All 0.5Gy data were well fitted with only three variables: two parameters for G_1_ and G_2_M CpMs (a weak block probability and a recycling rate in gen0,1,2, the same for G_1_ and G_2_M) and one for S phase delay (Fig. 10 lower left panel). G_1_ and G_2_M CpMs adopted for fitting 5Gy data were characterized by strong block probabilities but different recycling rates (lower in G_1_ than in G_2_M) and death rates (present only in G_1_ and higher than recycling rate) with a total of six variables. Similar results were obtained with D_R+CpM model (not shown). Actually, a correct prediction of the passage from one generation to the next requires models using a proper distribution of cell cycle duration (Fig. 8) and could be obtained with C_E (or D_R) models also without CpM (not shown). However, the models without CpM did not fit properly cell cycle percentages (not shown) and thus only C_E+CpM (or D_R+CpM) enabled contemporary satisfactory fit of both %gen and %G_1_/%S/G_2_M at all doses. The fact that this can be achieved without increasing the number of variables add support to the preference for these models.

## Notes

### Competing Interest Statement

The authors have declared no competing interest.

